# Why phosphenes are not always dots: modeling the shapes elicited by intracortical stimulation of human V1

**DOI:** 10.64898/2026.10.18.759925

**Authors:** Leili Soo, Eirini Schoinas, Geoffrey M. Boynton, Ione Fine, Eduardo Fernandez

**Affiliations:** Institute of Bioengineering, University Miguel Hernández, Elche, Spain; Department of Psychology, University of Washington, Seattle, USA; School of Biomedical Sciences, University of Leeds, Leeds, UK

**Keywords:** cortical visual prosthesis, phosphenes, primary visual cortex, intracortical microstimulation, computational model

## Abstract

The majority of visual cortical prosthetic devices currently implanted in patients or under development rely on small depth electrodes. Reports from patients implanted with these devices suggest that stimulation with tiny electrodes does not reliably produce small punctate spots, with some patients reporting a variety of percepts, including larger spots and irregularly shaped phosphenes. Here, we compare drawings and descriptions made by a cortical implant recipient to predicted phosphene patterns, generated using a “virtual patient” simulation based on a simple model of V1. We find that this model successfully reproduces the wide variety of percepts reported by this patient and further suggests that, for a subset of electrodes, stimulation may activate multiple locations — possibly as a result of axonal activation.

## 1 Introduction

Visual cortical prostheses aim to restore a limited but functionally useful form of vision in blind individuals by electrically stimulating neurons in primary visual cortex (V1), producing the subjective experience of small spots or patches of light known as phosphenes. The earliest characterization of cortical phosphenes comes from Brindley and Lewin’s 80-electrode chronic pial array [1], which reported that most individual phosphenes appeared as “a single very small spot of white light at a constant position.” However, a subset of electrodes reliably produced two or more spots, or “a small cloud,” rather than a single point. In a later paper, Rushton and Brindley [2] reported that some electrodes evoked markedly elongated, oval or bar-shaped percepts. Dobelle and colleagues noted that surface stimulation could, for some electrodes, produce clusters of several phosphenes [3, 4]. More recent work using subdural strip electrodes in both sighted (acute, pre-surgical) and blind (chronic, Orion-implanted) subjects found that most phosphenes resembled dots or blobs of 0.5-8 degrees in diameter [5, 6], though in one blind subject a subset of electrodes consistently produced short lines and a textured “oval area full of pinwheels” [7]. Like Rushton and Brindley, a few electrodes occasionally produced double phosphenes, which they attributed to the electrode lying across a sulcus.

The smaller number of intracortical microstimulation studies present a more heterogeneous picture. The earliest acute recordings, obtained intraoperatively in sighted patients during epilepsy surgery [8], found that single-electrode stimulation generally evoked a small, circular phosphene that was strongly colored near threshold — though one electrode produced a percept described as “an oblique line, narrow, blue.” Multi-electrode stimulation produced more complex percepts: two electrodes 0.7–1.0 mm apart produced irregular percepts described as “two blobs fusing” or “like a teddy bear,” and pairing two individually punctate “tiny, white dot” sources 0.9 mm apart yielded not two overlapping spots but a single elongated line. In a subsequent semi-chronic follow-up with 30–40 penetrating microwires [9], single-electrode stimulation near threshold generally evoked one small, round percept, ranging in apparent size from a “pin-point” to a “nickel” held at arm’s length, though some phosphenes were described as “fuzzy,” “amorphous,” or resembling “a drop of water”. Above threshold, percepts again became more complex with some electrodes producing multiple phosphenes.

In previous work using the same 96-channel Utah-array implant [10] as used in this paper, single-electrode stimulation mainly produced isolated, spatially localized elementary phosphenes, but these single-electrode phosphene shapes were heterogeneous across individuals: while one participant’s percepts were reliably punctate, for two other participants (including the patient of this paper) electrodes regularly produced non-punctate, structured shapes that were stable and reproducible within a given electrode across sessions. Users of the ICVP prosthesis [11] similarly report surprisingly complex shapes in a subset of electrodes. Together, these findings indicate that penetrating microelectrodes are capable of eliciting surprisingly complex phosphenes.

Despite this extensive collection of descriptive studies, it is not yet clear when and why complex shapes are sometimes but not always observed. The leftmost column of Figure 1 shows example patient drawings of complex phosphenes drawn by a CORTVIS participant intracortically implanted with a 96 electrode Utah array (the implant and surgical procedures are described previously [12]). The top five rows contain exemplars of drawings that were relatively common. Out of 64 electrodes 16 produced circular blobs, 14 produced rectangular percepts, 10 produced circular blobs that contained structure, 8 produced oriented lines, and 6 produced circular blobs with one or two intersecting lines. The bottom two rows show two examples of unique, complex percepts, which were found on 8 electrodes. Two electrodes produced oval blobs (not shown).

**Figure 1.**
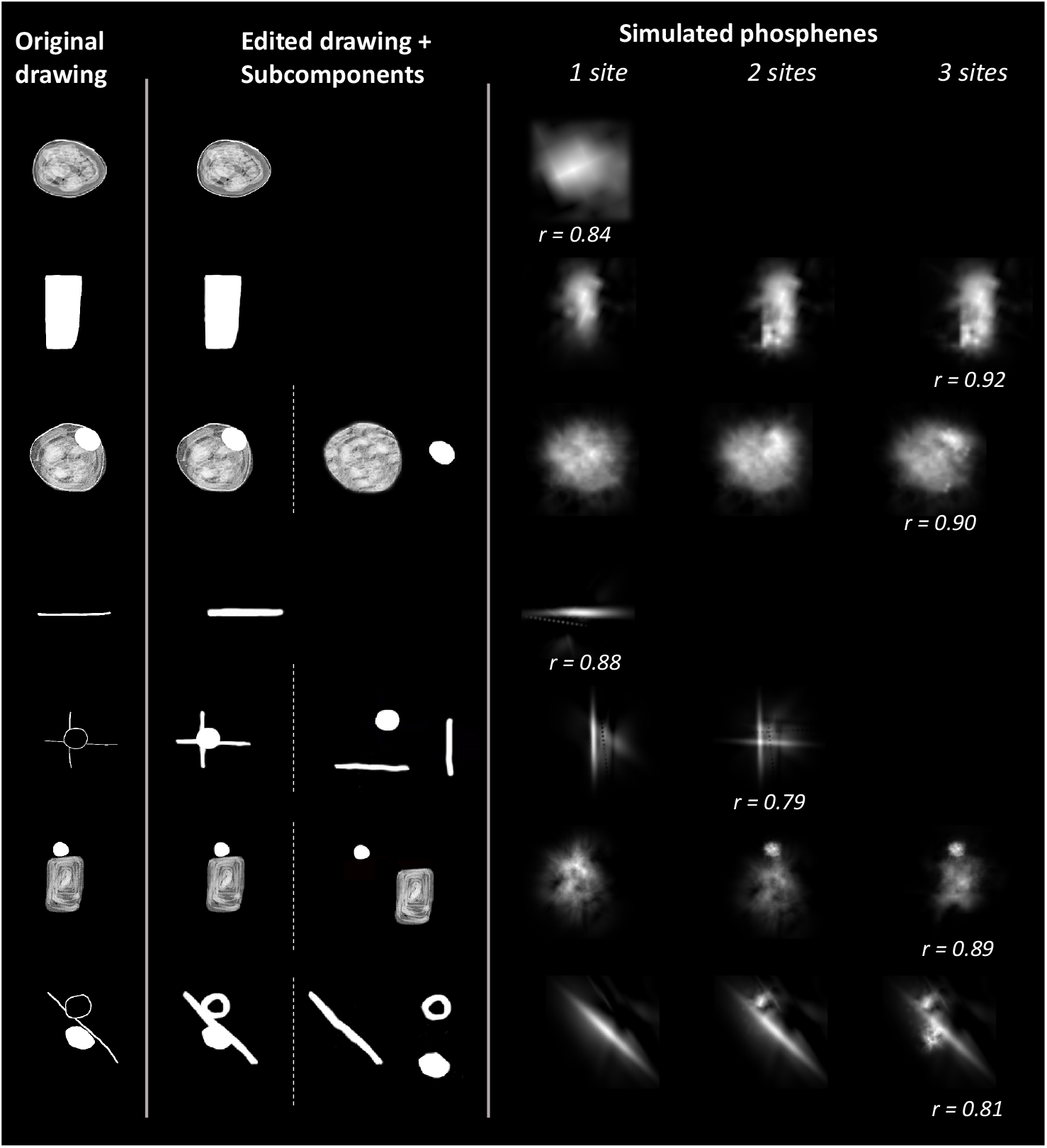
Simulated phosphenes provide a close match to patient drawings. The leftmost column shows the original patient drawings. The center column shows the edited images used to find closely matching phosphenes (Modeling Step I). The rightward column shows the best fitting combination of phosphenes elicited by 1, 2 or 3 electrode sites. The estimated number of sites was determined based on a sampling simulation, as described in Methods Step IV.

The center column shows edited drawings and sub-components (Modeling Step I) and the right column shows predicted phosphenes, as generated by our model. For each electrode we determined whether the phosphene was best fit by 1, 2 or 3 stimulation sites (Modeling Step IV). Where the phosphene was best fit with more than one stimulation site, the phosphenes produced by a smaller number of sites are shown.

How are such a variety of percepts generated? One possible cause of complex patient percepts is local stimulation of neurons tuned to multiple orientations as a result of an electrode lying on a discontinuity in the orientation pinwheel map. Figure 2 illustrates this with a simulation based on [13] representing a cortical region ∼7 mm from the foveal pole (∼0.4° eccentricity). Electrode A falls on an orientation pinwheel discontinuity, and therefore activates neurons tuned to both 30° and 120°, producing a complex percept.

**Figure 2.**
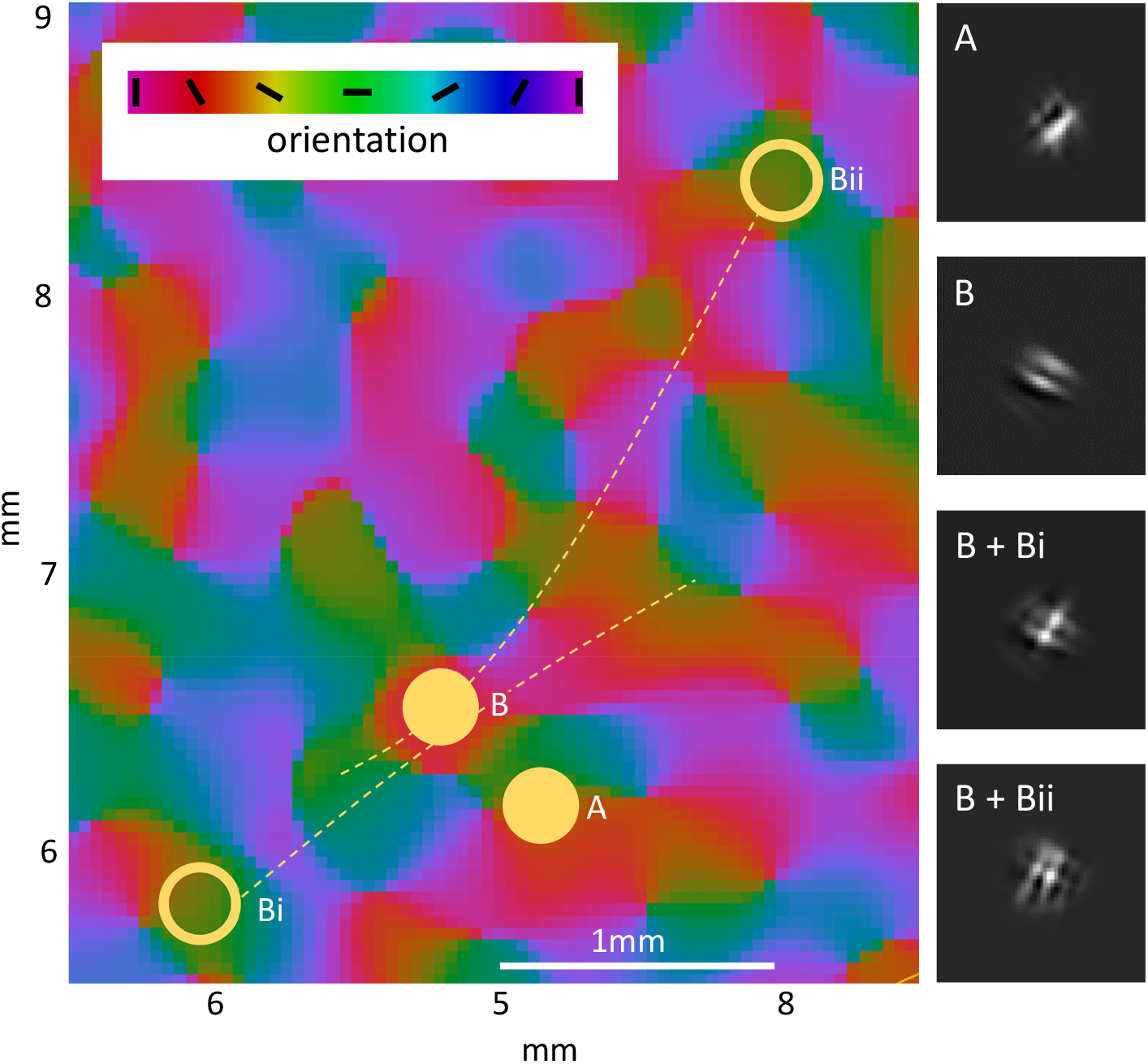
Both local and distal stimulation can result in complex percepts. Electrode A is located on a pinwheel discontinuity, and locally stimulates neurons tuned for both 30 and 120 degrees. Electrode B directly stimulates neurons primarily tuned to 30 degrees, but also stimulates passing axons from two regions, *Bi* and *Bii*, containing neurons tuned for 120 degrees. Right panels: virtual model predictions of elicited phosphenes. *Bi* shows the prediction when the distal stimulation is 1.0mm distant from the electrode, and *Bii* shows the prediction when the distal stimulation is 2.5mm distant. Even at a distance of 2.5mm the phosphene generated through axonal stimulation largely overlaps with the phosphene generated by local stimulation.

A second possible cause of complex percepts is stimulation of distal sites, due to stimulating axons of passage travelling under the electrode. Neurons lying directly under *ElectrodeB* are consistently tuned for 30 degrees, so direct local stimulation produces a pair of oriented bars. However, suppose that this electrode also stimulated passing axon fibers? Two examples of axonal fibers, both connected to regions tuned to 120 degrees, are illustrated by dashed lines. Percepts elicited bya local stimulation *B* in combination with *Bi* or *Bii* are considerably more complex. Note that even though *Bii* is 2.5 mm distant from *B* the phosphenes from the two sites still overlap.

Indeed, stimulated sites need to be surprisingly far apart to result in resolvable phosphenes. Figure 3 shows simulated phosphenes as a function of the separation between the activated cortical sites and eccentricity. At 0.5 degrees eccentricity it is not until stimulation sites are separated by ∼6.5mm that two sites produce clearly distinct phosphenes. (Because receptive fields vary linearly with eccentricity [14] and cortical magnification varies logarithmically [15] this ’resolvable’ distance varies across the cortical surface. At 1 degree eccentricity, sites must be separated by ∼ 5 mm, and at 4 degrees eccentric they would need to be separated by ∼4 mm [13].) Thus, the existence of non-overlapping phosphenes provides strong evidence for non-local stimulation from surprisingly distant locations in cortex. In Figure 1 the bottom two rows show spatially separated distinct phosphenes, but this was relatively rare, occurring in only 4/64 electrodes.

**Figure 3.**
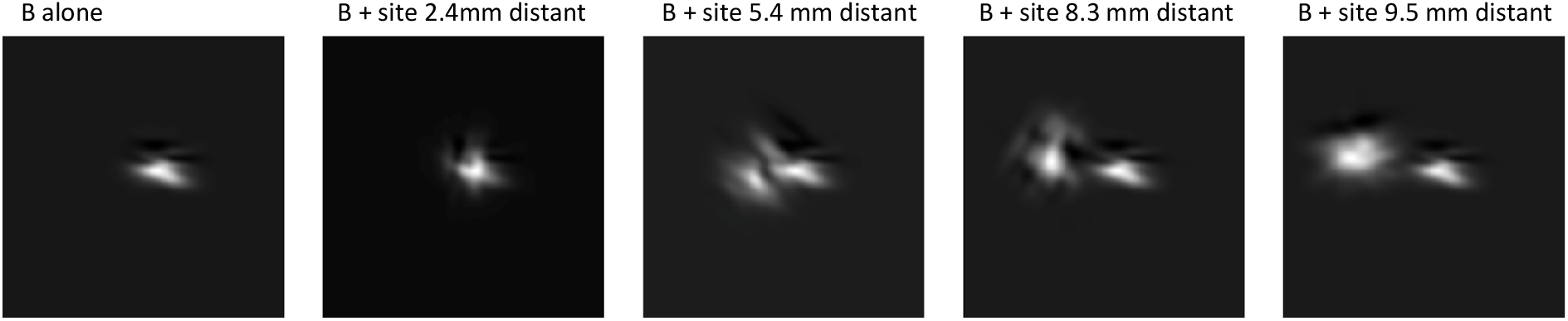
Stimulation sites need to be separated by a substantial distance before distinct phosphenes are elicited at the two sites.

Qualitatively, it seems that the neurophysiological architecture of early visual cortex is capable of generating surprisingly complex percepts. Our goal was to see whether this patient’s more complex percepts could be recreated using a simple model [13] that predicts the perceptual effects of cortical electrical stimulation, based on the neurophysiological architecture of V1. The model incorporates human cortical magnification [15, 16], orientation selectivity [17], ocular dominance [18], receptive field size [19], and the on and off structure of simple and complex neurons [20, 21]. Predicted percepts are generated by a linear sum of each neuron’s receptive field, weighted by the strength of electrical stimulation. This model has previously been shown to successfully predict percept brightness and size across a wide variety of studies. Here we evaluated whether it was also capable of predicting the diversity of patient-reported percepts, and whether model simulations were able to predict which percepts were plausibly elicited from local stimulation alone and which likely included activation of distal sites, as a result of stimulating passing axon fibers.

We compared each patient drawing to simulations based on our cortical model of V1. After generating 1000 random percepts, we selected the 24 percepts - *basis images* - that were most strongly correlated with either the edited patient drawing or a sub-component of that patient drawing, center column of Figure 1. We then found the best combination of up to 3 basis images, right column of Figure 1, that best predicted the patient-reported percept. We found that the model was remarkably successful at producing predicted phosphenes that qualitatively matched patient drawings.

To determine how many basis functions (sites) were required to best match the patient drawings we used a sampling simulation that statistically compared the increase in the correlation between the predicted phosphene and the edited patient drawing when the best-fitting basis image was added, as compared to adding a random phosphene image. We found that a subset of phosphenes were best fit by assuming multiple simulation sites, however some surprisingly complex percepts could be predicted from local stimulation alone.

Previously, we showed that our simple cortical model could quantitatively predict patient psychophysics. Here we show that although it is impossible to predict the percept that will be produced by an individual electrode, it is possible to predict the range of percepts likely to be experienced by patients.

## 2 Methods

### 2.1 Participants

The study was approved by the Clinical Research Committee of the General University Hospital of Elche and registered in ClinicalTrials.gov (NCT02983370). All relevant ethical guidelines, including those for clinical trials (EU No. 536/2014, repealed Directive 2001/20/EC), the Declaration of Helsinki, and European Commission Directives (2005/28/EC and 2003/94/EC), were strictly adhered to. All participants provided written informed consent before any procedures were performed and were regularly reminded of their rights throughout the experiment. Drawings and verbal reports were collected in a CORTVIS participant intracortically implanted with a 96 electrode Utah array capable of both stimulation and recording [12]). Reference and ground electrodes were uninsulated platinum wires positioned in brain tissue near the implanted array. The array was implanted in a location likely to include V1 and V2.

### 2.2 Patient Drawings

Methods are described fully in [22]. The participant drew the phosphene on a tablet mounted on a board at a comfortable distance. Before each stimulation, the participant was asked to maintain fixation on a tactile indicator glued to the middle of the tablet while their right index finger touched that fixation marker. The participant was informed that this represented the middle of their visual field and was instructed to reproduce the percept as if viewed at arm’s length. To capture relative brightness within the percept, participants shaded brighter regions with a higher density of pen strokes.

The drawings were then processed offline: first, the sketch was converted into a grayscale image and smoothed using a Gaussian filter to interpolate stroke density and reconstruct the perceived brightness gradients. Next, a threshold was applied to remove sparsely shaded artifacts generated when the stylus briefly contacted the tablet unintentionally; this step also sharpened edges that had been softened during Gaussian smoothing. Some additional corrections were made based on discrepancies between the drawing and the patient report. For example, if the patient reported a filled-in circular phosphene but there was a gap in the circular drawing, then the drawing was corrected to match. Finally, the cleaned image was rendered using a white color map to enhance percept contrast for display and analysis.

### 2.3 Modeling

#### 2.3.1 The pulse2percept virtual patient

The model was based on [23], where it is described in detail. Electric field spread was modeled based on the current-distance equation, 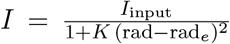 , where *I* is the current in *µ*A at a given location on the cortical surface, *I*_input_ is the stimulating current, and *rad*_*e*_ is the radial distance between that region of the cortical surface and the nearest region of the electrode [24]. Ocular dominance columns, orientation pinwheels and receptive fields were simulated based on Rojer and Schwartz [25]. Orientation columns were modeled by bandpass filtering white noise in the complex domain, with the resulting angle representing orientation preference. We modeled ocular dominance columns as the gradient of the same filtered white noise along a single direction, thereby generating orthogonal ocular dominance and orientation columns that closely resemble measured ocular dominance and orientation pinwheel maps as measured in humans [26]. The same band-pass filtered white noise that was used to generate orientation and ocular dominance maps was also used to generate the maps governing the separation and relative strength of receptive ON-and OFF-fields after bandpass filtering at twice the frequency used to generate orientation and ocular dominance columns [27].

Individual receptive fields were generated using a simple model that additively combined on and off sub-units with orientation, ocular dominance, and on-and off field organization drawn from the maps above[20]. The size of on and off subunits, and the spatial separation between them scaled linearly with eccentricity [19]. We assumed that the contribution of on neurons was weighted more heavily than the contribution of off neurons (*ω*_on-off_ = 0.9) enabling us to capture the phenomenon that phosphenes are occasionally dark at threshold, but are consistently bright as current increases above threshold. Predicted phosphenes were generated as a linear sum of receptive field profiles at each cortical location, weighted by the current stimulation intensity at that location.

In the original paper we fixed receptive field aspect ratios at 0.25 (such that a vertically oriented receptive field was always 4*x* as tall as it was wide). We replaced this with a distribution of receptive field aspect ratios based on distributions found in the macaque [28], such that the aspect ratios of receptive fields were drawn from a uniform distribution between 0.1 (extremely elongated) and 1

(circular). No other modifications to the model were made.

#### 2.3.2 Simulating phosphenes

Simulating phosphenes was a four-step process.

##### I CREATING OPTIMIZED PATIENT IMAGES

Each patient drawing (leftmost column of Figure 1) was edited manually to try to optimize featural similarity between the best-fitting phosphene images and patient images. For example, if a patient drew a thin line this would be thickened, to ensure that our fitting procedure captured this salient feature (middle column of Figure 1). For more complex patient drawings, as shown in the second column of Figure 1, we generated up to 3 sub-component patient images which allowed the model to separately fit individual features.

##### II SIMULATING INDIVIDUAL PHOSPHENES

We simulated 1000 phosphenes based on electrodes ranging in radius from 0.05 to 3mm microns (exponentially distributed so that small electrodes were common and large electrodes were rare) and eccentricities exponentially distributed between 0-15 degrees. For each electrode we generated a phosphene and then found the shift, rotation and scaling (since patient drawings were in arbitrary units) that produced the strongest correlation with either the manually edited patient image or one of its sub-components.

We saved the 24 phosphene basis images that were most strongly correlated with either the edited patient drawings or one of it’s subcomponents. If the original image had not been divided into sub-components, then the 24 basis images that were most strongly correlated with the patient image were saved. If the original patient drawing was divided into three sub-components (as was the case in the top row of Figure 1), then the 6 basis images that were most strongly correlated with the original image were saved, and 6 basis images were saved for each of the three sub-component images, for a total of 4 x 6 = 24 saved basis images. A final constraint ensured that saved basis images had pairwise correlations of *r <* 0.9, to avoid saving multiple almost identical basis images.

##### III. COMBINING PHOSPHENES

For each possible combination of basis images we used linear regression to find the weights that produced the best match to each edited phosphene, by minimizing ||*Xw* − *P* ||, where, *P* represents the edited patient image and is of size *n* x 1. *X* represents the basis images to be combined and is of size *n* x *c*, where *n* is the number of pixels in the image and *c* = 2 or 3, depending on how many basis images were combined.

For each possible combination of basis images we saved the correlation of *Xw* with the original edited patient image, *P* . This resulted in a total of 2324 possible combinations: 24 single basis images + 276 pairs + 2024 combinations of 3 basis images. For *c* = 1, 2 and 3 the basis image or combination of basis images (*X*) with the highest correlation was selected as the ’winning phosphene’.

##### IV. DETERMINING THE NUMBER OF STIMULATION SITES

We used a sampling simulation to determine whether improvements in correlation as a result of adding an additional optimized basis function were larger than the improvement in correlation that would be obtained by adding a random basis function. For each patient drawing, 24 random phosphene images were generated, using the same electrode size and cortical locations as for the saved basis functions of Step II, but with a freshly regenerated (random) cortical map for each. For each simulated phosphene we found the shift and scaling (but not rotation) that produced the strongest correlation with either the manually edited patient image or one of its sub-components. This produced a set of 24 random basis functions, each corresponding to one of the 24 optimized (selected from 1000 electrodes) basis functions of Step II.

To test whether the improvement in fit at each value of *c* exceeded what would be expected by chance, we constructed a matched null distribution. For each *c* we drew a single random phosphene basis function — not selected or optimized in any way — and combined it with every possible set of *c* − 1 optimized basis functions selected from Step II (excluding the basis function that was replaced), to produce a matrix *X*_*r*_ of size *n* x *c*. Regression weights were fit for each candidate *X*_*r*_ , and the combination with the highest correlation to *P* was retained. This was repeated over *R* = 100 iterations, each drawing a fresh random basis function and a freshly regenerated cortical map, to build a null distribution of *rX*_*r*_,*P* for each *c*. When *c* = 1, *X*_*r*_ reduces to a single random phosphene shifted and scaled to *P* . Our criterion was that if the real basis image outperformed at least (95%) of the null samples then adding that site improved the fit more than would be expected by chance.

## 3 Results

Figure 4 shows histograms comparing *rX,P* (the correlation between the best-fitting basis image or combination of basis images) to *rX*_*r*_,*P* (the correlations obtained when one of the basis images was replaced with a random basis image).

**Figure 4.**
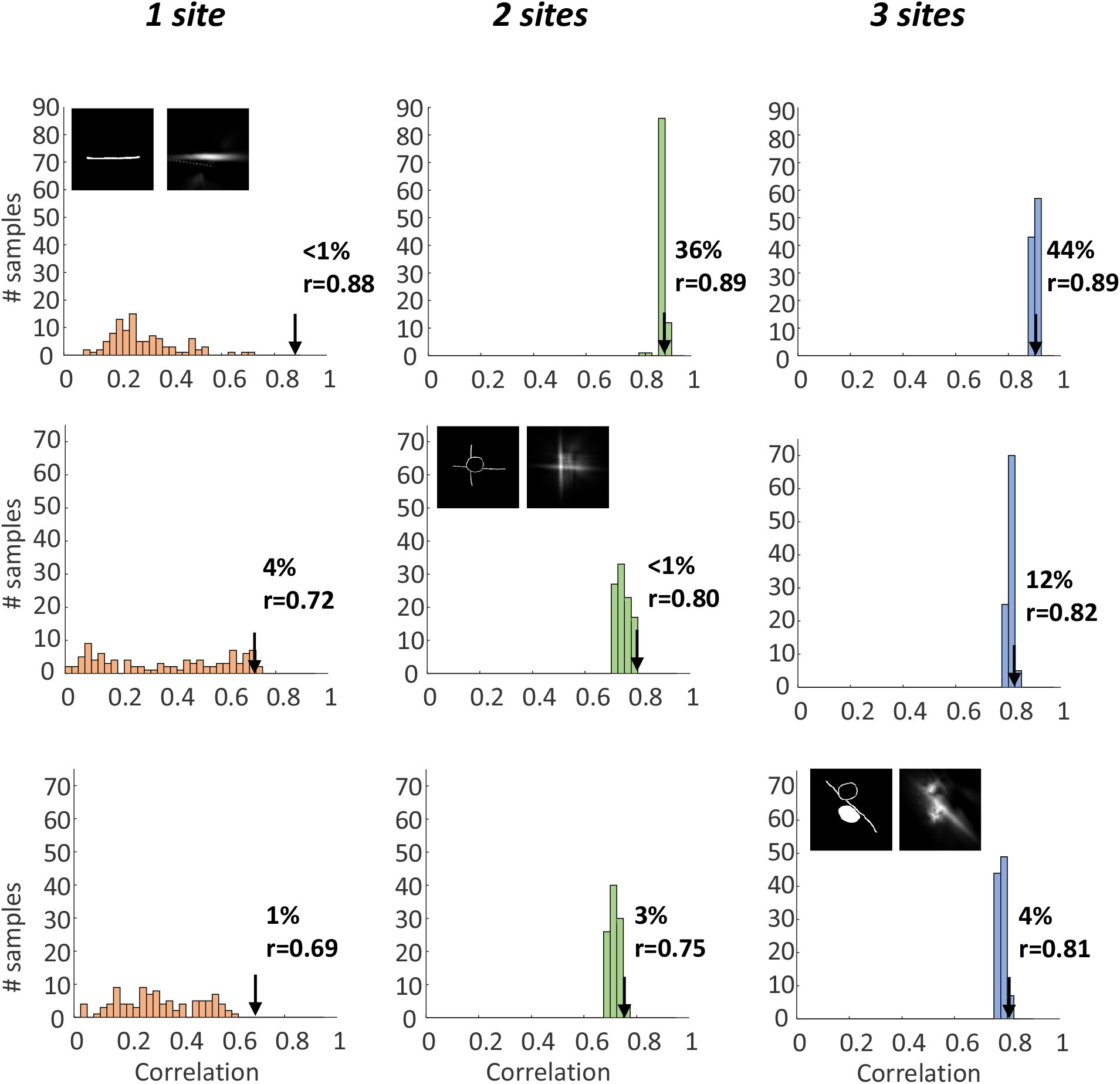
Sampling simulation used to determine the number of stimulated sites. Each row shows one patient drawing; columns show fits with 1, 2 or 3 basis images (sites). Histograms show the null distribution of correlations, *rX*_*r*_,*P* (100 repetitions), obtained when one of the basis images is replaced by a random, nonoptimized phosphene. The arrow marks the correlation obtained with the best-fitting real basis images *r*_*X,P*_ , and the value beside it gives that correlation (*r*). The percentage is the proportion of null samples with a correlation equal to or higher than the real fit. Insets show the edited patient drawing (left) and the best-fitting predicted phosphene (right) for the number of sites selected by the sampling simulation.

As expected, correlations always increase as a function of the number of basis images. This is both due to having more basis functions (i.e. free parameters in the regression fit to the patient drawing) and because, as the number of basis images increases, so too does the number of combinations from which *X* and *X*_*r*_ are selected on the basis of their correlation with the patient image.

Figure 5 summarizes the correlation between phosphene predictions and patient drawings. Correlations were high overall (median *r* = 0.89, range 0.41–0.96). A single site was sufficient for 13 of the 64 drawings (20%); for 21 drawings (33%) the best fit required two sites, and for 29 drawings (45%) three sites were required. One drawing was not fit significantly better than chance at any number of sites.

**Figure 5.**
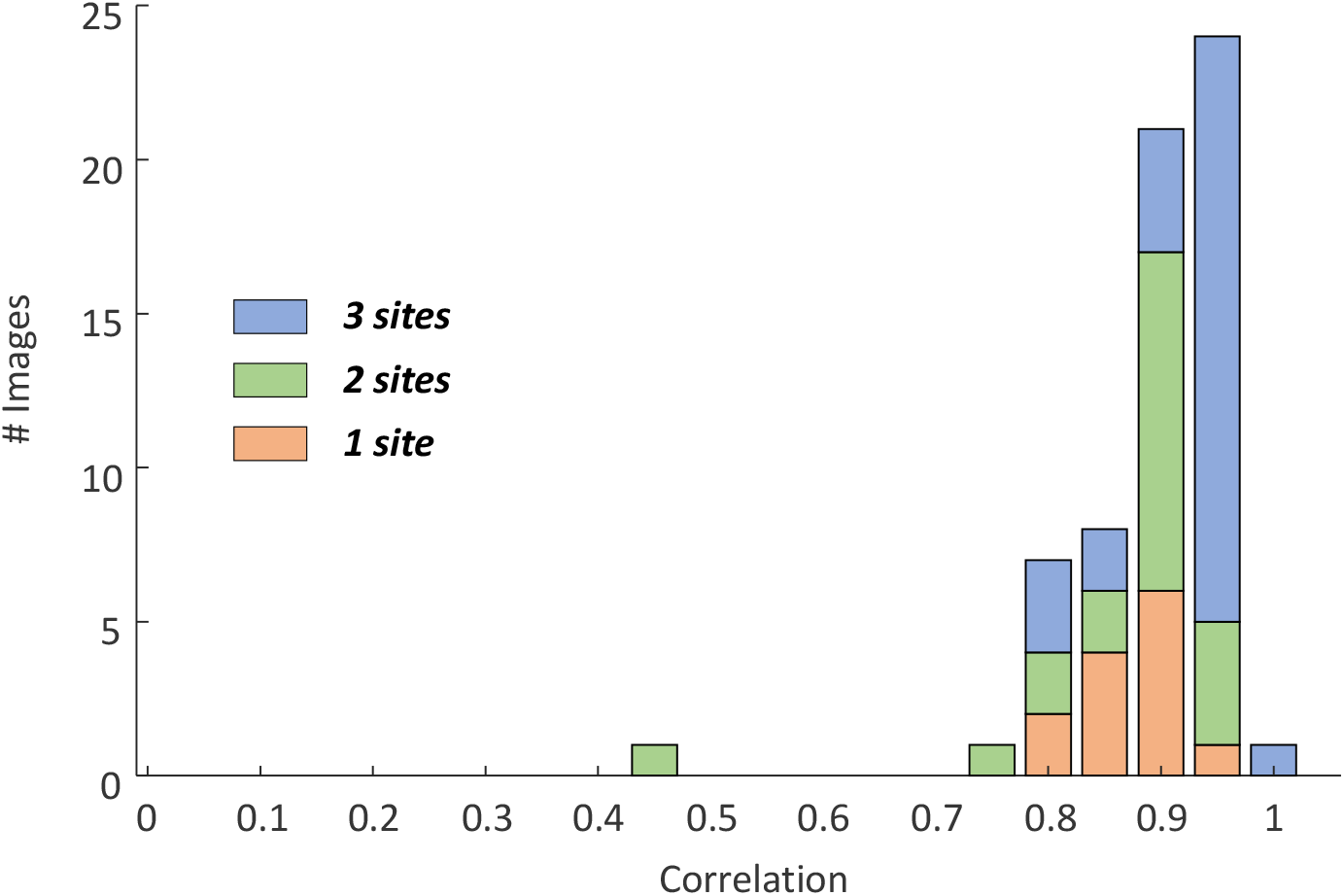
Fit quality. Stacked histogram of the correlation between the best-fitting predicted phosphene and the edited patient drawing (*r*_*X,P*_). Colors indicate the number of predicted sites (1, 2 or 3) based on our sampling simulation.

## 4 Discussion

One important issue that cortical prosthetic devices will need to compensate for is the variability in phosphene shape across different electrodes. This variability seems not to be due to measurement noise. As described in previous work [22], phosphene identity was stable and reliably discriminable across repeated testing, suggesting that the idiosyncratic appearance of phosphenes is largely a fixed, reproducible property of a given stimulation site rather than the result of trial-to-trial randomness.

We show here that a simple model of cortex — incorporating cortical magnification, receptive field size and location, orientation pinwheel and ocular dominance maps, and on/off subfield structure with an on-center bias, summed linearly and weighted by current spread — reproduced much of the diversity elicited by small intracortical depth electrodes. Because small electrodes stimulate only a handful of cortical cells, the resulting phosphene reflects the specific local mix of receptive field properties at that site; irregular, non-punctate phosphenes are an expected consequence of stimulating small, spatially localized cortical territory.

When we included axonal stimulation in our model the complexity of predicted phosphenes increased. For over two thirds of the electrodes we found that fits to patient percepts were significantly better when multiple basis functions (sites) were included, suggesting axonal activation as a plausible mechanism for explaining some of the more idiosyncratic percepts seen in this patient. At one level, it is reassuring that the bizarre, complex percepts reported by some — but not all — patients are not mysterious or idiosyncratic to a particular device or patient. They are a direct consequence of fine-grained cortical architecture — pinwheels, ocular dominance columns, on/off structure, and axonal geometry — and this kind of variability can be modeled and predicted *in general*. What our model cannot do is predict, for a given patient, what the phosphene elicited by a given *one specific* electrode will look like in advance: that depends on the precise local arrangement of cortical structure at that electrode’s exact location.

In the retinal domain, a promising proposed approach has been to treat each electrode’s own phosphene as a fixed perceptual “basis function” and to build complex percepts by combining these basis functions [29]. However, in cortex these basis functions cannot be predicted from retinal geometry; the phosphene produced by a given electrode must be measured empirically, electrode by electrode. It is difficult to see how this would scale to high-density arrays. Current devices do include recording capability, so in principle complex receptive fields could be characterized via multi-unit recordings rather than behavioral report — but this is unlikely to be straightforward in practice: electrode position and responsiveness drift over time, patients have limited tolerance for lengthy mapping sessions, and it remains unclear how well receptive fields estimated this way would correspond to what patients actually perceive.

The major challenge for cortical sight recovery is not engineering. Building denser, more stable, better-recording arrays, difficult as that is, is likely to be tractable. ’Hacking’ the neural code of early visual cortex is a substantially harder problem — one that may prove intractable using our current concepts and tools.

## Acknowledgments

Claude (multiple versions) was used for help in looking for code bugs, commenting code, and proof reading.

## Author Roles

L.S.: Conceptualization, Methodology, Investigation, Data Curation, Writing. E.S.: Conceptualization, Software, Visualization, Formal Analysis, Writing, G.M.B: Software, Writing. I.F.: Conceptualization, Methodology, Software, Visualization, Writing, Funding Acquisition. E.F. Conceptualization, Methodology, Writing, Funding Acquisition.

## References

[1] Brindley G S and Lewin W S 1968 The sensations produced by electrical stimulation of the visual cortex J Physiol 196 479–93 URL http://www.ncbi.nlm.nih.gov/entrez/query.fcgi?cmd=Retrieve&db=PubMed&dopt=Citation&list_uids=4871047

[2] Rushton D N and Brindley G S 1978 Properties of cortical electrical phosphenes Frontiers in Visual Science ed Cool S J and Smith E L (Berlin, Heidelberg: Springer Berlin Heidelberg) pp 574–593 ISBN 978-3-540-35397-3

[3] Dobelle W H and Mladejovsky M G 1974 Phosphenes produced by electrical stimulation of human occipital cortex, and their application to the development of a prosthesis for the blind J Physiol 243 553–76 URL http://www.ncbi.nlm.nih.gov/entrez/query.fcgi?cmd=Retrieve&db=PubMed&dopt=Citation&list_uids=4449074

[4] Dobelle W H, Mladejovsky M G, Evans J R, Roberts T S and Girvin J P 1976 Braille reading by a blind volunteer by visual cortex stimulation Nature 259 111–112 doi: 10.1038/259111a0

[5] Girvin J, Evans J, Dobelle W, Mladejovsky M, Henderson D, Abramov I, Gordon J and Turkel J 1979 Electrical stimulation of human visual cortex: the effect of stimulus parameters on phosphene threshold Sensory Processes 3 66–81 ISSN 0363-3799

[6] Bosking W H, Sun P, Ozker M, Pei X, Foster B L, Beauchamp M S and Yoshor D 2017 Saturation in phosphene size with increasing current levels delivered to human visual cortex J Neurosci 37 7188–7197 ISSN 1529-2401 (Electronic) 0270-6474 (Print) 0270-6474 (Linking) doi: 10.1523/JNEUROSCI.2896-16.2017

[7] Niketeghad S, Muralidharan A, Patel U, Dorn J D, Bonelli L, Greenberg R J and Pouratian N 2020 Phosphene perceptions and safety of chronic visual cortex stimulation in a blind subject Journal of Neurosurgery JNS 132 2000–2007 doi: 10.3171/2019.3.JNS182774

[8] Bak M, Girvin J P, Hambrecht F T, Kufta C V, Loeb G E and Schmidt E M 1990 Visual sensations produced by intracortical microstimulation of the human occipital cortex Medical & Biological Engineering & Computing 28 257–259 doi: 10.1007/BF02442682

[9] Schmidt E M, Bak M J, Hambrecht F T, Kufta C V, Rourke D K O and Vallabhanath P 1996 Feasibility of a visual prosthesis for the blind based on intracortical microstimulation of the visual cortex Brain 119 507–522

[10] Fernández E, Alfaro A, Soto-Sánchez C, Gonzalez-Lopez P, Lozano A M, Peña S, Grima M D, Rodil A, Gómez B, Chen X, Roelfsema P R, Rolston J D, Davis T S and Normann R A 2021 Visual percepts evoked with an intracortical 96-channel microelectrode array inserted in human occipital cortex J Clin Invest 131 ISSN 1558-8238 (Electronic) 0021-9738 (Print) 0021-9738 (Linking) doi: 10.1172/jci151331

[11] Troyk P R 2017 The intracortical visual prosthesis project Artificial Vision: A Clinical Guide ed Gabel V P (Cham: Springer International Publishing) pp 203–214 doi: 10.1007/978-3-319-41876-6_16

[12] Grani F, Soto-Sanchez C, Farfan F D, Alfaro A, Grima M D, Rodil Doblado A and Fernández E 2022 Time stability and connectivity analysis with an intracortical 96-channel microelectrode array inserted in human visual cortex J Neural Eng 19 ISSN 1741-2552 doi: 0.1088/1741-2552/ac801d

[13] Fine I and Boynton G M 2015 Pulse trains to percepts: the challenge of creating a perceptually intelligible world with sight recovery technologies Philosophical Transactions of the Royal Society B: Biological Sciences 370 20140208 ISSN 0962-8436

[14] Freeman J and Simoncelli E P 2011 Metamers of the ventral stream Nat Neurosci 14 1195–201 ISSN 1546-1726 (Electronic) 1097-6256 (Print) 1097-6256 (Linking) doi: 10.1038/nn.2889

[15] Schwartz E L 1980 Computational anatomy and functional architecture of striate cortex: a spatial mapping approach to perceptual coding Vision Res 20 645–69 URL http://www.ncbi.nlm.nih.gov/entrez/query.fcgi?cmd=Retrieve&db=PubMed&dopt=Citation&list_uids=7445436

[16] Schwartz E L 1980 A quantitative model of the functional architecture of human striate cortex with application to visual illusion and cortical texture analysis Biol Cybern 37 63–76 URL http://www.ncbi.nlm.nih.gov/htbin-post/Entrez/query?db=m&form=6&dopt=r&uid=6772241

[17] Obermayer K and Blasdel G G 1993 Geometry of orientation and ocular dominance columns in monkey striate cortex J Neurosci 13 4114–29

[18] Adams D L, Sincich L C and Horton J C 2007 Complete pattern of ocular dominance columns in human primary visual cortex J Neurosci 27 10391–403 ISSN 1529-2401 (Electronic) 0270-6474 (Print) 0270-6474 (Linking) doi: 10.1523/JNEUROSCI.2923-07.2007

[19] Keliris G A, Li Q, Papanikolaou A, Logothetis N K and Smirnakis S M 2019 Estimating average single-neuron visual receptive field sizes by fmri Proc Natl Acad Sci U S A 116 6425–6434 ISSN 1091-6490 (Electronic) 0027-8424 (Print) 0027-8424 (Linking) doi: 10.1073/pnas.1809612116

[20] Mata M L and Ringach D L 2005 Spatial overlap of on and off subregions and its relation to response modulation ratio in macaque primary visual cortex J Neurophysiol 93 919–28 ISSN 0022-3077 (Print) 0022-3077 (Linking) doi: 10.1152/jn.00668.2004

[21] Najafian S, Koch E, Teh K L, Jin J, Rahimi-Nasrabadi H, Zaidi Q, Kremkow J and Alonso J M 2022 A theory of cortical map formation in the visual brain Nature Communications 13 2303 ISSN 2041-1723 doi: 10.1038/s41467-022-29433-y

[22] Soo L, Rodil Doblado A, Grani F, Lopez Peco R, Villamarin-Ortiz A, Morollon Ruiz R, Waclawczyk D, Fine I, Gonzalez-Lopez P, Soto-Sanchez C, Alfaro A and Fernandez E 2026 Intracortical microstimulation evokes diverse electrode-specific phosphenes in the early visual cortex of blind individuals medRxiv Preprint doi: 10.64898/2026.10.05.26364412

[23] Fine I and Boynton G M 2024 A virtual patient simulation modeling the neural and perceptual effects of human visual cortical stimulation, from pulse trains to percepts Scientific Reports 14 17400 doi: 10.1038/s41598-024-65337-1

[24] Tehovnik E J and Slocum W M 2007 Phosphene induction by microstimulation of macaque v1 Brain Res Rev 53 337–43 ISSN 0165-0173 (Print) 0165-0173 (Linking) doi: 10.1016/j.brainresrev.2006.11.001

[25] Rojer A S and Schwartz E L 1990 Cat and monkey cortical columnar patterns modeled by bandpass-filtered 2d white noise Biol Cybern 62 381–91 ISSN 0340-1200 (Print) 0340-1200 (Linking)

[26] Yacoub E, Harel N and Ugurbil K 2008 High-field fmri unveils orientation columns in humans Proc Natl Acad Sci U S A 105 10607–12 ISSN 1091-6490 (Electronic) 0027-8424 (Print) 0027-8424 (Linking) doi: 10.1073/pnas.0804110105

[27] Rahimi-Nasrabadi H, Jin J, Mazade R, Pons C, Najafian S and Alonso J M 2021 Image luminance changes contrast sensitivity in visual cortex Cell Rep 34 108692 ISSN 2211-1247 (Electronic) doi: 10.1016/j.celrep.2021.108692

[28] Ringach D L 2002 Spatial structure and symmetry of simple-cell receptive fields in macaque primary visual cortex J Neurophysiol 88 455–63 ISSN 0022-3077 (Print) 0022-3077 (Linking) doi: 10.1152/jn.2002.88.1.455

[29] Granley J, Relic L and Beyeler M 2022 Hybrid neural autoencoders for stimulus encoding in visual and other sensory neuroprostheses Advances in Neural Information Processing Systems vol 35 pp 22671–22685 doi: 10.52202/068431-1647

